# Species-Specific Formation of Paraspeckles in Intestinal Epithelium Revealed by Characterization of *NEAT1* in Naked Mole-rat

**DOI:** 10.1101/2022.02.17.480918

**Authors:** Akihiro Yamada, Hikaru Toya, Mayuko Tanahashi, Misuzu Kurihara, Mari Mito, Shintaro Iwasaki, Satoshi Kurosaka, Toru Takumi, Archa Fox, Yoshimi Kawamura, Kyoko Miura, Shinichi Nakagawa

**Affiliations:** RNA Biology Laboratory, Faculty of Pharmaceutical Sciences, Hokkaido University, Kita 12-jo Nishi 6-chome, Kita-ku, Sapporo 060-0812, Japan; RNA Systems Biochemistry Laboratory, RIKEN Cluster for Pioneering Research, 2-1 Hirosawa, Wako, Saitama 351-0198, Japan; Department of Computational Biology and Medical Sciences, Graduate School of Frontier Sciences, The University of Tokyo, 5-1-5 Kashiwanoha, Kashiwa, Chiba 277-8561, Japan; RIKEN Brain Science Institute, 2-1 Hirosawa, Wako, Saitama 351-0198, Japan; Department of Physiology and Cell Biology, Kobe University School of Medicine, 7-5-1 Kusunoki-cho, Chuo, Kobe 670-0017, Japan; School of Human Sciences, University of Western Australia, Crawley, Western Australia, 6009; Harry Perkins Institute of Medical Research, Nedlands, Western Australia, Australia; School of Molecular Sciences, University of Western Australia, Crawley, Western Australia, 6009, Australia; Department of Aging and Longevity Research, Faculty of Life Sciences, Kumamoto University, 1-1-1 Honjo, Chuo-ku, Kumamoto 860-8556, Japan; Center for Metabolic Regulation of Healthy Aging, Kumamoto University, Kumamoto 860-8556, Japan

## Abstract

Paraspeckles are mammalian-specific nuclear bodies built on the long noncoding RNA *NEAT1_2*. The molecular mechanisms of paraspeckle formation have been mainly studied using human or mouse cells, and it is not known if the same molecular components are involved in the formation of paraspeckles in other mammalian species. We thus investigated the expression pattern of *NEAT1_2* in naked mole-rats (n*NEAT1_2*), which exhibit extreme longevity and lower susceptibility to cancer. In the intestine, n*NEAT1_2* is widely expressed along the entire intestinal epithelium, which is different from the expression of m*Neat1_2*, that is restricted to the cells of the distal tip in mice. Notably, the expression of FUS, a FET family RNA binding protein, essential for the formation of paraspeckles both in humans and mice, was absent in the distal part of the intestinal epithelium in naked mole-rats. Instead, mRNAs of other FET family proteins EWSR1 and TAF15 were expressed in the distal region. Exogenous expression of these proteins in Fus-deficient murine embryonic fibroblast cells rescued the formation of paraspeckles. These observations suggest that n*NEAT1_2* recruits different set of RNA binding proteins in a cell type-specific manner during the formation of paraspeckles in different organisms.

## Introduction

A large number of non-coding RNAs are transcribed from the genomes of higher eukaryotes, constituting a significant portion of the transcriptional output (Clark et al. 2011; Djebali et al. 2012). Noncoding RNA transcripts longer than 200 nucleotides are collectively called long noncoding RNAs (lncRNAs), and a series of studies revealed that they are functionally divided into several groups (reviewed in Kopp and Mendell 2018; Ali and Grote 2020; Gil and Ulitsky 2020; Nakagawa et al. 2021; Statello et al. 2021). One group of lncRNAs are epigenetic regulators that control gene expression via association with chromatin modifying complexes *in cis* or *trans* (reviewed in Ali and Grote 2020; Gil and Ulitsky 2020; Statello et al. 2021). Others function as molecular sponges that sequester associating RNA binding proteins or microRNAs and negatively regulate their function (reviewed in Kopp and Mendell 2018). Architectural RNAs (arcRNAs) are an emerging group of lncRNAs that serve as a structural component of submicron-scale nonmembranous organelles, which are often referred to as molecular condensates (reviewed in Nakagawa et al. 2021).

*MALAT1* and *NEAT1* are highly abundant lncRNAs, which are located at the same syntenic chromosomal regions in humans and mice (Hutchinson et al. 2007). *MALAT1* is localized to nuclear speckles that are enriched in splicing regulators such as SR proteins and UsnRNPs. While *MALAT1* is not required for the formation of nuclear speckles (Eissmann et al. 2012; Nakagawa et al. 2012; Zhang et al. 2012), it has been proposed to regulate pre-mRNA splicing via association with splicing regulators (Tripathi et al. 2010). *NEAT1* is the most extensively studied arcRNA that plays essential role to build mammalian-specific nuclear bodies called paraspeckles (reviewed in Fox et al. 2018; Nakagawa et al. 2018), which were originally defined as nuclear bodies enriched in the RNA binding protein PSPC1 and 2 (paraspeckle protein 1 and 2) (Fox et al. 2002). Two isoforms of *NEAT1, NEAT1_1* and *NEAT1_2* are transcribed from the same promoter, and the longer isoform *NEAT1_2* plays the architectural role whereas *NEAT1_1* alone cannot induce the formation of paraspeckles (Sasaki et al. 2009; Naganuma et al. 2012; Li et al. 2017; Isobe et al. 2020). Paraspeckles contain more than 30 RNA binding proteins (Naganuma et al. 2012; Fong et al. 2013), which are radially arranged to form spheres with characteristic core-shell structure (Souquere et al. 2010; West et al. 2016). A group of paraspeckle proteins including NONO and FUS are indispensable for the formation of paraspeckles (Naganuma et al. 2012; West et al. 2016). Many of these essential paraspeckle proteins contain distinct intrinsically disordered regions or prion-like domains, which undergo phase transition *in vitro* to form a hydrogel (Hennig et al. 2015). *NEAT1* induces the formation of phase-separated condensates by recruiting paraspeckle proteins onto distinct regions of its transcript (Yamazaki et al. 2018). Paraspeckle spheres fuse to form sausage-or tube-like structures with distinct radial diameter, suggesting that they behave as flexible block co-polymer micelles rather than rigid complexes with components strictly located at specific positions (Yamazaki et al. 2021).

Paraspeckles are ubiquitously observed in most cultured cell lines (reviewed in Fox et al. 2018; Nakagawa et al. 2018), with a few exceptions such as ES cells that do not express enough *NEAT1_2* (Modic et al. 2019). On the other hand, prominent expressions of *Neat1_2* in mice (m*Neat1_2*) is restricted to a subset of particular cell types, including distal tip cells of intestinal epithelium and corpus luteal cells in female ovaries (Nakagawa et al. 2011; Isobe et al. 2020). m*Neat1* knockout mice exhibit various phenotypes including decreased fertility and impaired mammary tract formation in female animals (Nakagawa et al. 2014; Standaert et al. 2014); however, the penetrance of the sub-fertile phenotype is around 50% and is variable even in the same individuals (Nakagawa et al. 2014). In addition, the lack of m*Neat1* results in an increase or decrease in tumorigenesis in different tumor models, although the expression of m*Neat1* is under the control of p53 pathway in both cases (Adriaens et al. 2016; Mello et al. 2017). These observations suggest that m*Neat1* plays a physiological role in a condition-or context-dependent manner in different cell types or experimental models (reviewed in Nakagawa et al. 2018).

Given *Neat1* expression and paraspeckle formation in tissues of living animals have been mostly studied in mice, it is possible that the molecular components of paraspeckles that associate with *Neat1* could vary between different mammalian species. To investigate the molecular conservation of paraspeckle components between species, here we studied the expression pattern of *NEAT1* in naked mole-rats (n*NEAT1*), an animal characterized by longevity and low susceptibility to cancer formation (reviewed in Buffenstein et al. 2022). We found characteristic expression of n*NEAT1_2* in the intestinal epithelium of naked mole-rats, however this was observed throughout the entire length of the intestinal epithelium, unlike the restricted expression in mice. In addition, nFUS, an essential RNA binding protein needed to form paraspeckles, was absent in the distal part of intestinal epithelium in naked mole-rats. Instead, mRNAs of n*EWSR1* and n*TAF15* were broadly expressed throughout the epithelium, suggesting that the FET family proteins substitute for the function of nFUS to form paraspeckles in the distal regions of intestinal epithelium in naked mole-rats. These observations suggest that paraspeckle components are variable between species, raising a possibility of species-specific function of paraspeckles in certain tissues or cell types.

## Results

### Identification of *MALAT1* and *NEAT1* in naked mole-rats

To get insights into the role of lncRNA in naked mole-rats, we initially tried to identify *MALAT1* and *NEAT1*, two of the most abundant lncRNAs, whose genes are adjacently located in the human and mouse genome (Hutchinson et al. 2007). We performed RNA-Seq analyses using RNAs obtained from Naked mole-rat skin fibroblast cells (NS-Fibroblasts), and found that a large number of reads were mapped to two distinct regions in the assembly JH602080 of naked mole-rat genome (hetGla2), flanked by *SCYL1* and *FRMD8* (Figure 1A), which corresponds to a region syntenic to human chromosome 11 and mouse chromosome 19 containing the two lncRNAs and the same neighboring genes (Figure 1A). Sequences at the 3’ regions of naked mole-rat *MALAT1* and *NEAT1* (hereafter n*MALAT1* and n*NEAT1*) exhibited remarkable homology to mouse/human *MALAT1*/*NEAT1* (hereafter m*Malat1*, m*Neat1*, h*MALAT1*, h*NEAT1*, respectively) (Figure 1B), suggesting that n*MALAT1* and n*NEAT1* receive 3’ processing characteristics of these lncRNAs, which include the cleavage by RNaseP and the formation of triple-helix structures that stabilize the transcripts (Wilusz et al. 2008; Wilusz et al. 2012). To determine the transcription start sites of n*MALAT1* and n*NEAT1*, we performed 5’ RACE (<u>r</u>apid <u>a</u>mplification of <u>c</u>DNA <u>e</u>nds) PCR. The PCR fragments were cloned into a plasmid vector and we sequenced three clones for each gene, which yielded the same sequences shown in Figure 1C. Probes designed against n*MALAT1* and n*NEAT1* detected bands of the same size as m*Neat1* and m*Malat1* by Northern blot (Figure 1D, E), where the sizes were consistent with the aforementioned prediction. Probes that detect the 5’ region of n*NEAT1* detected two bands (Figure 1E), suggesting that naked mole-rat also have a short (n*NEAT1_1*) and a long (n*NEAT1_2*) isoform of n*NEAT1*, like humans and mice (Sasaki et al. 2009; Sunwoo et al. 2009).

**Figure 1.**
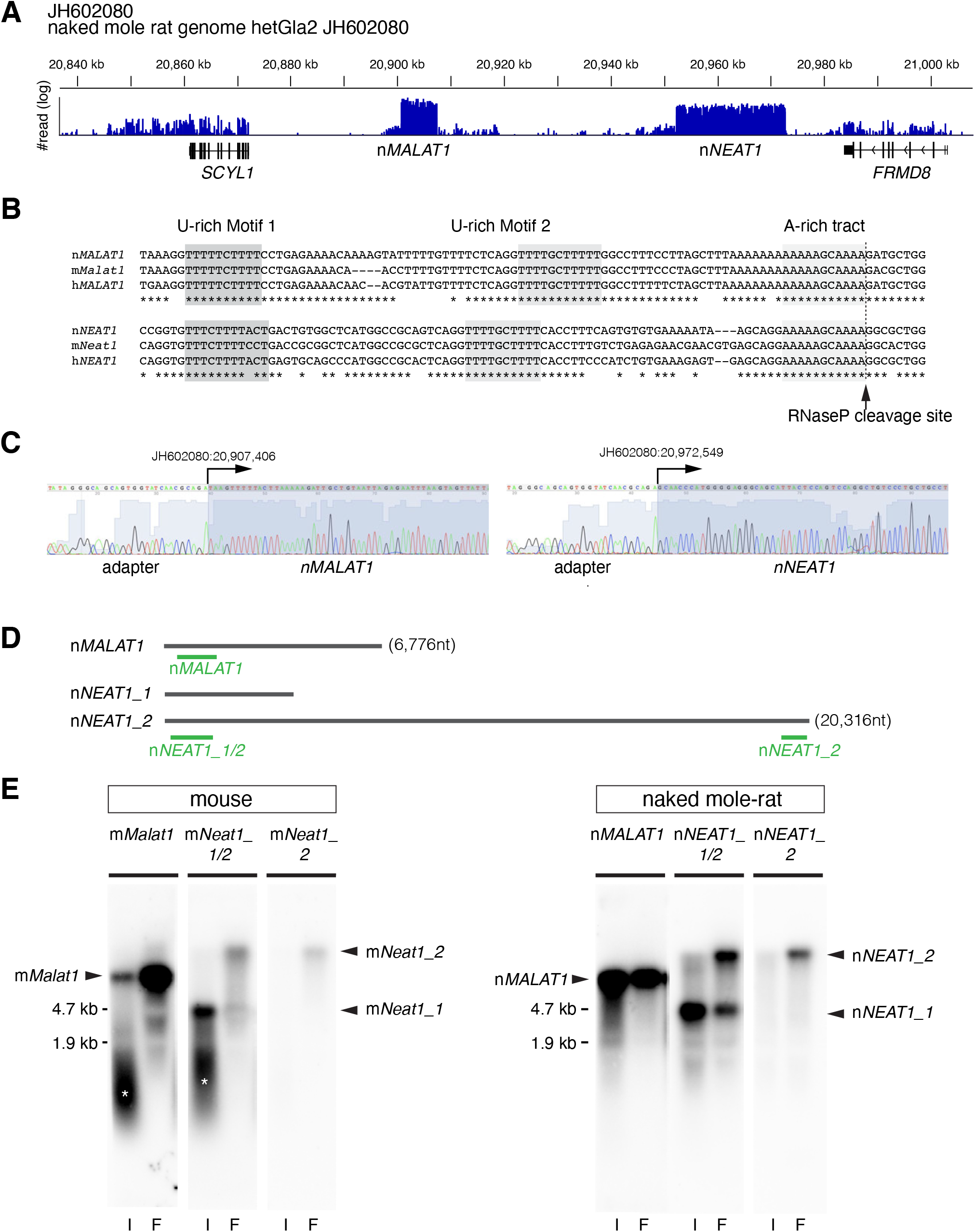
Identification of *MALAT1* and *NEAT1* in naked mole-rats. (A) Distribution of RNA-Seq reads mapped to the region of the naked mole-rat genome syntenic to mouse chromosome 19 containing m*Malat1* (mouse *Malat1*) and m*Neat1* (mouse *Neat1*), which are flanked by *SCYL1* and *FRMD8*. Note that two regions were abundantly transcribed into RNA, which corresponded to n*MALAT1* (naked mole-rat *MALAT1*) and n*NEAT1* (naked mole-rat *NEAT1*), respectively. (B) Sequence alignment of the 3’ regions of *NEAT1* and *MALAT1* in naked mole-rats (n), mouse (m), and human (h). The tandem U-rich motifs followed by A-rich tract, which form triple helix structures, are highlighted in shadow boxes and are highly conserved between the three species. The arrow indicates the putative RNaseP cleavage site that generate the 3’ end of these transcripts. (C) Sequence analyses of PCR fragments obtained by the 5’ RACE analyses. Arrows indicate the positions of the transcription start sites of n*MALAT1* and n*NEAT1*. (D) Schematic drawing indicating the positions of probes used for Northern blot and *in situ* hybridization studies. (E) Northern blot analyses of n*MALAT1* and n*NEAT1* expression in embryonic fibroblast cells and intestine. I: intestine, F: fibroblasts. Asterisks shows the signals derived from degraded RNAs.

To further characterize n*MALAT1* and n*NEAT1* in naked mole-rats, we examined the subcellular distribution of these transcripts in skin fibroblast cells (NS-Fibroblasts) (Figure 2). h*MALAT1*/m*Malat1* is localized in nuclear speckles that are enriched in splicing regulators (Hutchinson et al. 2007; Tripathi et al. 2010). The localization of n*MALAT1* coincided with nSRSF1, a marker for nuclear speckles in NS-Fibroblasts, which was indistinguishable from the localization of m*Malat1* in mouse embryonic fibroblast cells (MEF) (Figure 2A, Supplementary figure S1). To test if n*NEAT1* is localized in paraspeckles, we examined if a series of antibodies that recognize paraspeckle marker proteins in mice (West et al. 2016) also recognized antigens in naked mole-rats. However, only two of these antibodies, those that detect nNONO and nFUS were effective in naked mole-rats (A.Y. and S.N., unpublished observations), and both of these proteins are essential for the formation of paraspeckles in mice. n*NEAT1_2*, the architectural isoform of *NEAT1*, exhibited the same localization in paraspeckles visualized by marker protein nNONO and nFUS in NS-Fibroblasts, as was the case for the colocalization of m*Neat1_2* and these marker proteins in MEF (Figure 2B, Supplementary figure S1). To test if the paraspeckles in naked mole-rats possess the characteristic core-shell structures found in human or mouse paraspeckles (Souquere et al. 2010; West et al. 2016), we observed them in NS-Fibroblasts using structured illumination microscopy (Figure 2C, Supplementary figure S1). Simultaneous detection of n*NEAT1* and the paraspeckle markers nNONO and nFUS revealed that the 5’ and 3’ regions of n*NEAT1* were located at the peripheral regions of paraspeckles, whereas nNONO and nFUS were located in the core (Figure 2C, Supplementary figure S1), as was reported in previous studies in humans and mice (Souquere et al. 2010; West et al. 2016). Taken together, we concluded we have correctly identified homologues of *MALAT1* and *NEAT1* in naked mole-rats, which exhibited almost identical subcellular distributions compared to the mouse homologues.

**Figure 2.**
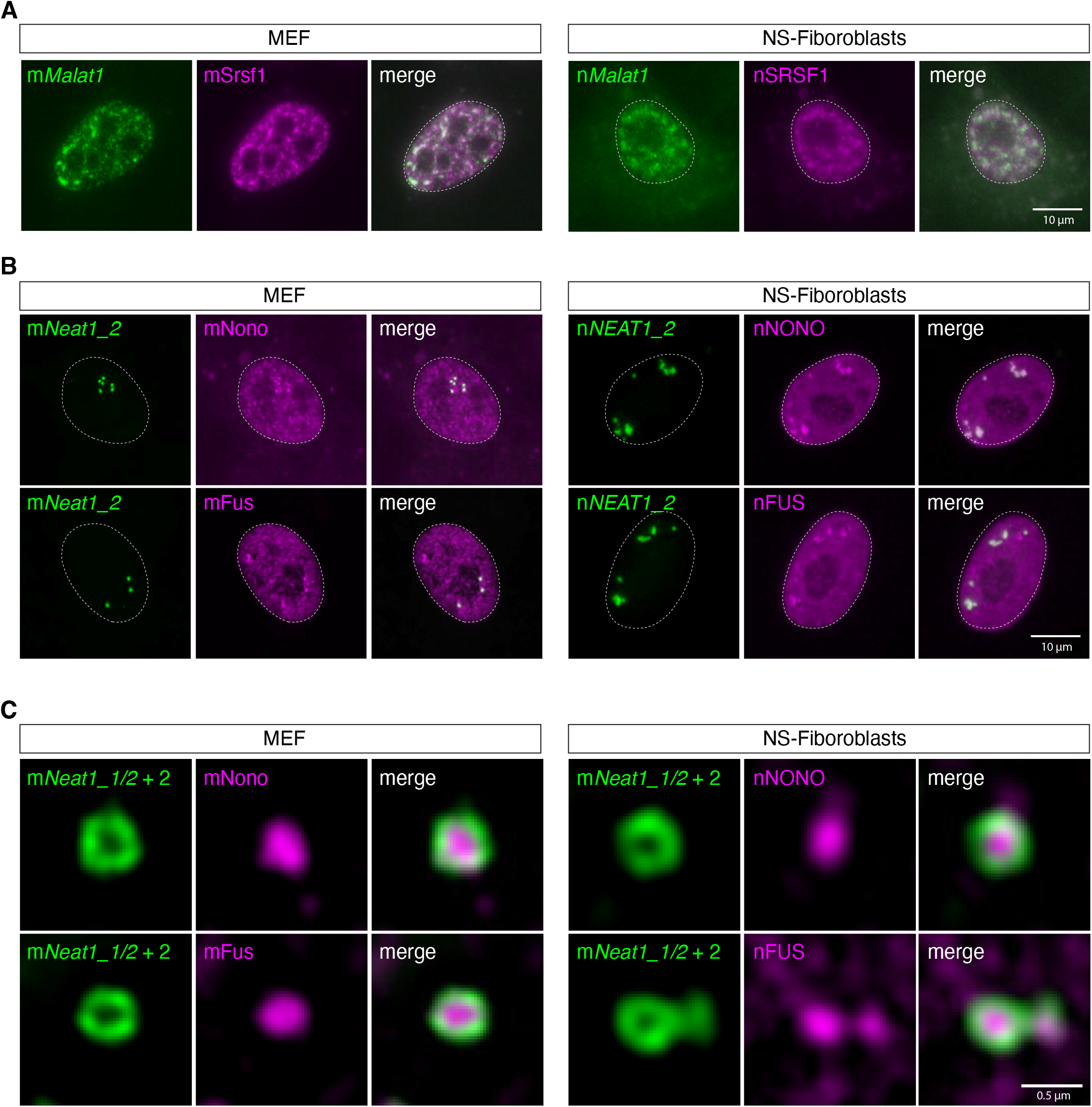
The same subcellular distribution of m*Malat1*/n*MALAT1* and m*Neat1*/n*NEAT1* in MEF and NS-Fibroblasts. (A) Simultaneous detection of m*Malat1*/n*MALAT1* (green) and the nuclear speckle marker Srsf1/SRSF1 (magenta) in MEF and NS-Fibroblasts. Note that both m*Malat1* and n*MALAT1* are enriched in nuclear speckles identified by mSrsf1/nSRSF1 expression. (B) Simultaneous detection of m*Neat1_2*/n*NEAT1_2* (green) and the paraspeckle markers mNono/mFus and nNONO/nFUS in MEF and NS-Fibroblasts, respectively. (C) Simultaneous detection of m*Neat1_2*/n*NEAT1_2* (green) and mNono/mFus and nNONO/nFUS (magenta) at high magnification using structural illumination microscopy. Scale bars, 10 µm in A and B and 0.5 µm in C.

### n*NEAT1_2* is broadly expressed in the intestinal epithelium to form paraspeckles in naked mole-rats

We then examined the expression pattern of n*MALAT1* and n*NEAT1* in the liver and intestine of naked mole-rats, two representative tissues that exhibit ubiquitous and cell type-specific expression of m*Neat1_2* in mice, respectively (Nakagawa et al. 2011; Isobe et al. 2020). n*MALAT1* was strongly expressed in all the cells in these tissues (Figure 3A), the same as the expression of m*Malat1* in mice (Nakagawa et al. 2012). n*NEAT1_1* also exhibited the same expression pattern as m*Neat1_1*, however, n*NEAT1_2* was broadly expressed along the entire length of the epithelium in naked mole-rats (right magenta boxes in Figure 3A and B). The expression pattern of n*NEAT1_2* was very different to that of m*Neat1_2*, which exhibited highly restricted expression, only in the distal tip cells of the intestinal epithelium (left magenta boxes in Figure 3A and B). The higher expression of n*NEAT1_2* in the intestinal epithelium compared to m*Neat1_2* was also confirmed by RT-qPCR (Figure 3C). Because paraspeckle formation is dependent on the expression of *NEAT1_2* (Sasaki et al. 2009; Naganuma et al. 2012; Li et al. 2017; Isobe et al. 2020), these observations suggested that paraspeckle formation in naked mole-rats is more broadly observed in the intestinal epithelium than in mice. To confirm this hypothesis, we investigated the colocalization of n*NEAT1_2* and the paraspeckle marker protein nNONO in the intestine of naked mole-rats (Figure 4A, Supplementary figure S1). As previously reported, colocalization of m*Neat1_2* and mNono was found only in the tip of the intestinal epithelium in mice (Figure 4A, Supplementary figure S1). On the other hand, n*NEAT1_2* signals overlapped with nNONO, both in the distal tip, and the bottom region of the intestinal epithelium (Figure 4A, Supplementary figure S1), suggesting that paraspeckle formation was rather ubiquitously observed in the intestine of naked mole-rats. These observations suggested that paraspeckle formation is differentially regulated by species-specific expression of *NEAT1_2* in particular tissue or cell types in each species.

**Figure 3.**
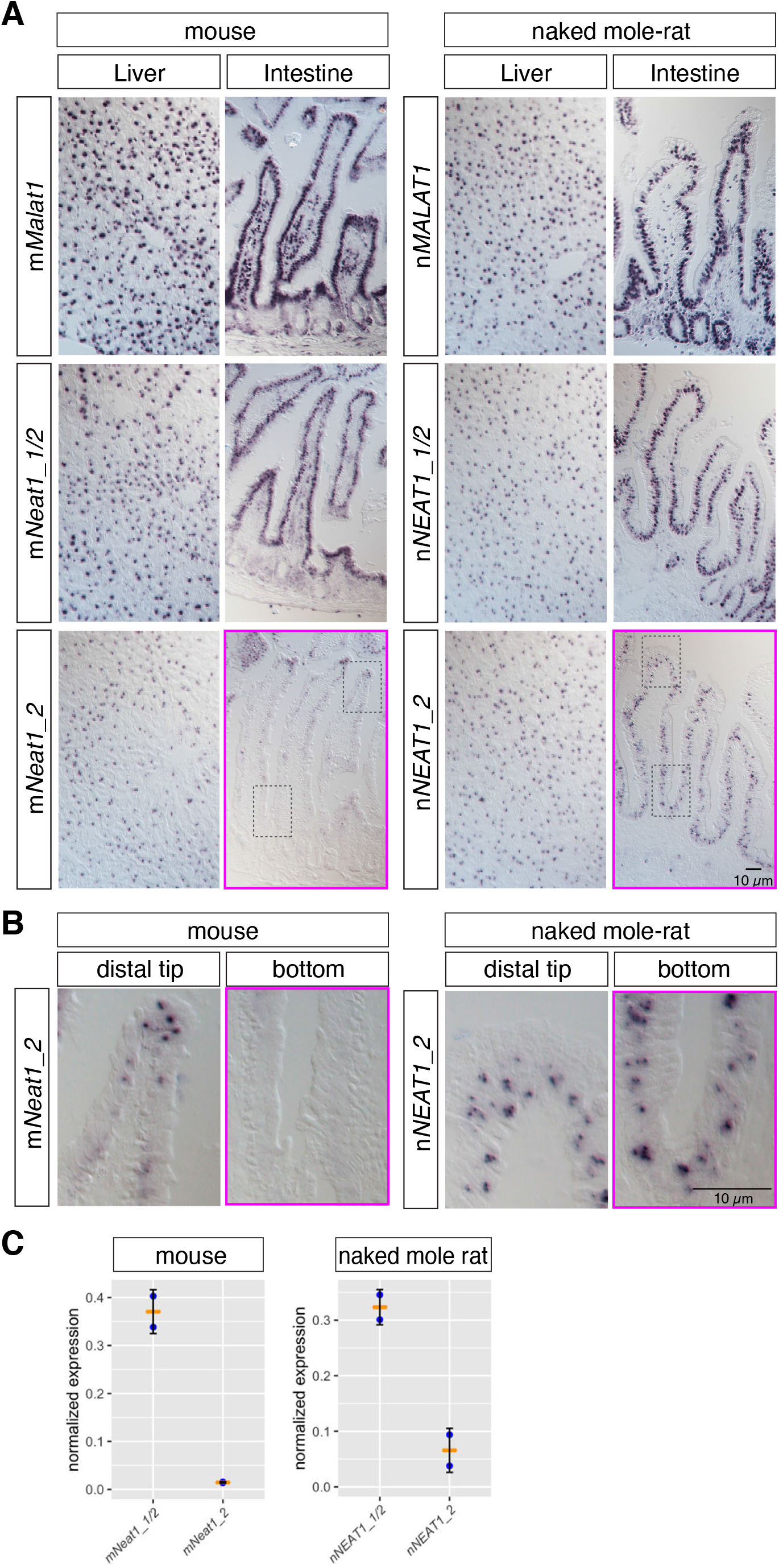
n*NEAT1_2* is expressed in the entire regions of intestinal epithelium in naked mole-rats. (A) *In situ* hybridization analyses of m*Malat1*/n*MALAT1* and m*Neat1*/n*NEAT1* in liver and intestine of mice and naked mole-rats. Note that the expression of m*Neat1_2* is restricted to the distal tip cells in the intestinal epithelium of mice, whereas the expression of n*NEAT1_2* is broadly observed throughout the epithelium in naked mole-rats (magenta boxes). (B) Higher magnification images of regions indicated in dashed boxes in A. Note that m*Neat1_2* is not expressed in the bottom region of intestinal epithelium. Scale bars, 10 µm. (C) RT-qPCR analyses of the expression of m*Neat1*/n*NEAT1* in intestine. The relative expression of n*NEAT1_2* to total n*NEAT1_1/2* is elevated compared to the ratio of m*Neat1_2* to total m*Neat1_1/2*.

**Figure 4.**
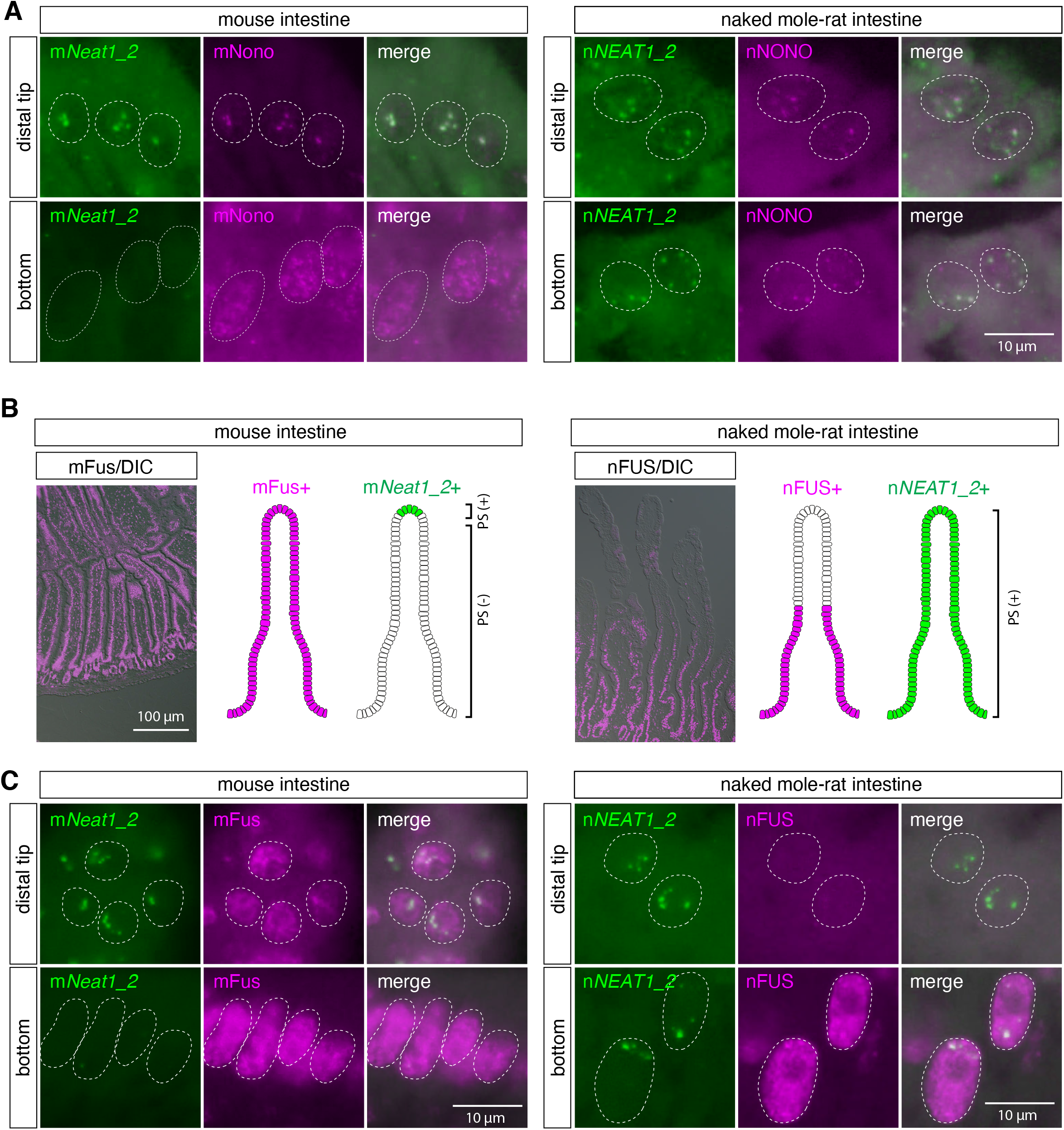
The essential paraspeckle component FUS is not expressed in the distal region of intestinal epithelium in naked mole-rats. (A) Simultaneous detection of m*Neat1_2*/n*NEAT1_2* (green) and the paraspeckle marker mNono/nNONO (magenta) in the top and the bottom regions of intestinal epithelium in mice and naked mole-rats. Paraspeckle formation was observed in the bottom region of intestinal epithelium in naked mole-rats. (B) The expression of mFus/nFUS (magenta) in intestinal epithelium in mice and naked mole-rats and schematic drawing of the expression of mFus/nFUS and m*Neat1_2*/n*NEAT1_2* (green) in the epithelium. DIC: differential interference contrast images. (C) Simultaneous detection m*Neat1_2*/n*NEAT1_2* (green), and mFus/nFUS (magenta) in the top and the bottom regions of intestinal epithelium in mice and naked mole-rats. Note the formation of paraspeckles in top region of the epithelium of naked mole-rats despite the lack of the expression of essential paraspeckle protein nFUS. Scale bars, 10 µm in A and C and 100 µm in B.

### The essential paraspeckle protein FUS is not expressed in the distal regions of intestinal epithelium despite the formation of paraspeckles in naked mole-rats

To further confirm the formation of paraspeckles in the basal region of intestinal epithelium in naked mole-rats, we examined the localization of nFUS, an essential component of paraspeckles that assembles the *NEAT1* ribonucleoprotein complex to build the nuclear bodies (Naganuma et al. 2012; Shelkovnikova et al. 2014; West et al. 2016). Unexpectedly, the expression of nFUS was not detected in the distal regions of intestinal epithelium in naked mole-rats (Figure 4B, C), despite the distinct formation of paraspeckles confirmed by the co-localization of n*NEAT1_2* and nNONO (Figure 4A, Supplementary figure S1). On the other hand, mFus was ubiquitously expressed in the entire region of intestinal epithelium in mice (Figure 4B, C). The lack of nFUS expression in the paraspeckle-forming cells was rather inconsistent with a previous observation showing that hFUS/mFus is essential for the formation of paraspeckles in human and mouse (Naganuma et al. 2012; Shelkovnikova et al. 2014; West et al. 2016).

To investigate the molecular mechanisms that lead to the formation of paraspeckles in the absence of nFUS in the distal region of intestinal epithelium in naked mole-rats, we examined the expression of n*EWSR1* and n*TAF15*, the two other members of the FET (FUS, EWS, and TAF15) protein family that share similar domain structures characterized by N-terminally located long prion-like domains (PrLD) followed by RNA recognition motifs (RRM), zf-RanBP domain, and the second PrLD containing RGG repeats (Schwartz et al. 2015) (Figure 5A, Supplementary figure S2A). The putative amino acid sequences of naked mole-rat FET family proteins were highly homologous to human or mouse genes, showing <95% identity at the amino acid level (Figure 5B, Supplementary figure S2B). Because antibodies that recognize nEWSR1 and nTAF15 in naked mole-rats were not available as far as we tested, we firstly performed RT-qPCR to examine the expression of each FET family protein in the intestine of mice and naked mole-rats (Figure 5C). We also estimated the expression of FET family proteins in MEF and NS-Fibroblasts, by RT-qPCR and the analyses of RNA-Seq data, respectively (Figure 5C). To quantitate the relative abundance of mRNA of each gene in total RNAs, we normalized the efficiency of PCR primers used for RT-qPCR analyses using 0.1 pmol of the PCR fragments. The normalized Ct values revealed that the expression of m*Fus* is much higher than m*Ewsr1* in mouse intestine, whereas the expression of n*EWSR1* was higher than n*FUS* in naked mole-rat intestine (Figure 5C). The same trend was also observed in fibroblast cells, and the expression of m*Fus* was higher than m*Ewsr1* in MEF, whereas the expression of n*EWSR1* was higher than n*FUS* in NS-Fibroblasts (Figure 5C). On the other hand, the expression of m*TAF15*/n*TAF15* was higher in intestine and lower in fibroblast cells both in mice and naked mole-rats (Figure 5C). We then examined the spatial expression pattern of mRNAs of FET family proteins in the intestine of mice and naked mole-rats using *in situ* hybridization. In mice, m*Fus* was widely expressed in the entire intestinal epithelium (Figure 5D, E). On the other hand, the expression of n*FUS* was restricted to the basal region of intestinal epithelium (Figure 5F, G), consistent with the lack of nFUS expression in the distal region of the epithelium in naked mole-rats (Figure 4B). These observations suggested that nEWSR1 and nTAF15 redundantly regulate the formation of paraspeckles in the distal tip cells that lack the expression of nFUS in naked mole-rats. The area expressing n*FUS* mRNA was smaller than the protein expression (Figure 4B), suggesting that nFUS expression lasted for a longer period in the cells that stopped the mRNA expression, which cells were displaced distally during the turnover of the epithelium.

**Figure 5.**
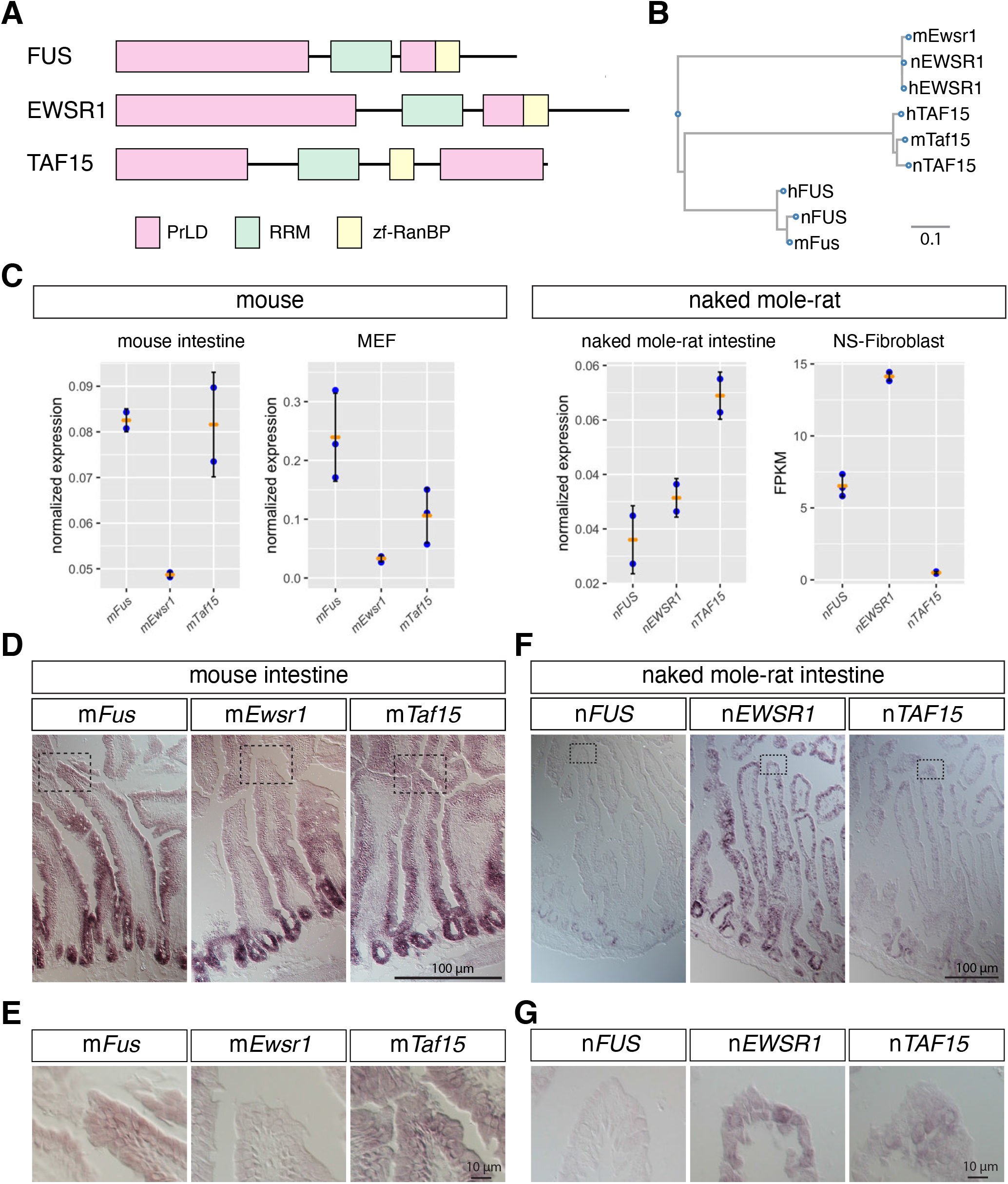
mRNAs of FET family proteins are redundantly expressed in intestine of naked mole-rats. (A) Domain structures of FET family proteins. PrLD: prion like domain, RRM: RNA recognition motif, zf-RanBP: zinc finger motif in Ran binding protein. The PrLD were predicted using PLAAC (Lancaster et al. 2014). (B) Phylogenetic tree of FET family proteins of human (h), mouse (m), and naked mole-rat (n). Scale bar indicates the amino acid substitution ratio per residue. (C) Quantification of gene expression of FET family proteins. The expression was estimated by normalized RT-qPCR analyses for intestine and MEF, and by RNA-Seq analyses for NS-Fibroblasts. FPKM: Fragments Per Kilobase of transcript per Million mapped reads. Note the higher expression of m*Fus* compared to m*EWSR*1 in mice, and the lower expression of n*FUS* compared to n*EWSR1* in naked mole-rats. (D-G) *In situ* hybridization analyses of the expression of FET family proteins in the intestine of mice (D, E) and naked mole-rats (F, G). Dashed boxes in D and F indicate the regions shown at a higher magnification in E and G. Note that n*EWSR1* and n*TAF15* were expressed in the distal tip cells of the intestinal epithelium of naked mole-rats. Scale bars, 100 µm in D and F, and 10 µm in E and G.

### FET family proteins redundantly regulate the formation of paraspeckles in MEF derived from m*Fus* knockout mice

Finally, we examined if mEwsr1 and mTaf15 can redundantly induce the formation of paraspeckles using MEF derived from m*Fus* knockout (KO) mice (West et al. 2016). Because only putative protein sequences were available for naked mole-rats (i.e., Refseq ID with suffix XP), we used FET family genes from mouse for the rescue experiments (Figure 6A). As we have previously reported (West et al. 2016), the exogenous expression of mFus restored the formation of paraspeckles, which was confirmed by co-localization of m*NEAT1_2* and mNono (Figure 6B). The formation of paraspeckles was also observed in the m*Fus* KO MEF expressing mEwsr1 and mTaf15 (Figure 6B), indicating that the FET family proteins redundantly function to regulate the formation of paraspeckles.

**Figure 6.**
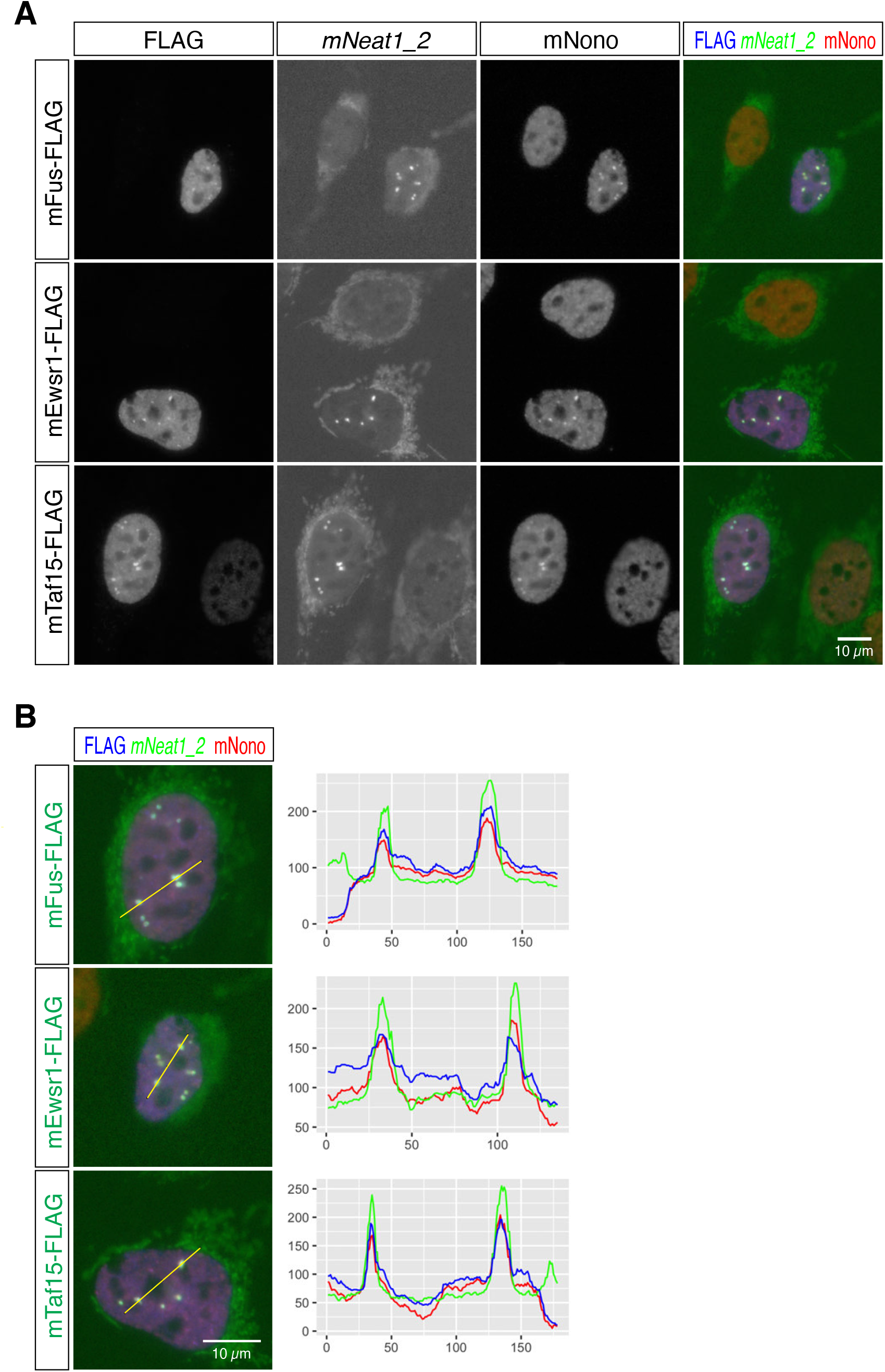
FET family proteins redundantly rescue the formation of paraspeckles in MEF lacking the expression of mFus. (A) Simultaneous detection of m*Neat1_2*, exogenously-expressed FLAG-tagged FET family proteins, and the paraspeckle marker mNono in MEF derived from m*Fus* KO mice. Note the absence of paraspeckle formation in m*Fus* KO MEF and the rescue of the paraspeckle formation in the cells expressing FET family proteins. The cytoplasmic signals in the green channel (m*Neat1_2* channel) were derived from cross-reactions of streptavidin to endogenous biotinylated proteins in the mitochondria. (B) Merged images of exogenously-expressed FLAG-tagged FET family proteins (blue), m*Neat1_2* (green), and paraspeckle marker mNono (red) and the intensity profile plots. Yellow lines indicate the region analyzed in the intensity profile plots. X axis represents the number of pixels and Y axis represents arbitrary unit for the intensity of the signals of each channel. Scale bars, 10 µm.

## Discussion

Here we have revealed the expression patterns and subcellular localizations of the homologues of the two abundant lncRNAs, n*MALAT1* and n*NEAT1*, in naked mole-rats. While the expression of n*MALAT1* was almost the same as the expression of m*Malat1*, we observed striking differences between the expression of n*NEAT1* and m*Neat1* in intestinal epithelium. n*NEAT1_2* was broadly expressed along the entire length of the epithelium, and paraspeckle formation was ubiquitously observed in the n*NEAT1_2* expressing cells. The expression pattern of n*NEAT1_2* is quite different from that of m*Neat1_2*, which is highly restricted to the distal tip cells of the intestinal epithelium in mice. On the other hand, the uniform expression of n*NEAT1_1*/m*Neat1_1* was commonly observed in both mice and naked mole-rats. The production of h*NEAT1_2* in HeLa cells is controlled by opposing activities of CPSF6 and HNRNPK that promotes and inhibits the use of the alternative polyadenylation site required for h*NEAT1_1* production, respectively (Naganuma et al. 2012). However, it remains unknown if a similar mechanism is operating in tissues of living animals, because the two RNA binding proteins are ubiquitously expressed in all of the intestinal epithelium in mice (S.N., unpublished observation), despite the variable ratios of the expression of m*Neat1_1* and m*Neat1_2* in different tissues and cell types (Isobe et al. 2020).

It should be noted that the half-life of p53 is 10 times longer in embryonic fibroblast cells from naked mole-rats when compared with human or mouse embryonic fibroblast cells, resulting in constitutive accumulation of p53 in the nucleus in the absence of DNA damage (Deuker et al. 2020). Considering m*Neat1* is regulated under the control of p53 (Adriaens et al. 2016; Mello et al. 2017), the increased amount of p53 possibly leads to hyper-activation of n*NEAT1* promoter in naked mole-rats. The overproduction of n*NEAT1* may induce preferential readthrough of 5’ located polyadenylation, resulting in the production of n*NEAT1_2* in a broader range of cells in the intestinal epithelium. It is also possible that species-specific modification of HNRNPK and/or CPSF6 may modulate the activity of these proteins, resulting in the differential use of the polyadenylation signal in mice and naked mole-rats. It will be essential to investigate the expression pattern of paraspeckle proteins and other regulatory proteins in detail to further clarify the molecular mechanisms that lead to rather ubiquitous formation of paraspeckles in the intestine of naked mole-rats.

Although the distal region of the intestinal epithelium lacked the expression of nFUS in naked mole-rats, distinct formation of paraspeckles was nevertheless observed there. This observation contradicted the prior observations from mice and cell lines that hFUS/mFus plays an essential role to build paraspeckles (Naganuma et al. 2012; Shelkovnikova et al. 2014; West et al. 2016). We have shown that two FET-family RNA binding proteins, n*EWSR1* and n*TAF15*, were expressed in the nFUS-negative distal regions, at least at the mRNA level, in naked mole-rats, and the expression of these proteins rescued the formation of paraspeckles in MEF lacking the expression of mFus. The formation of paraspeckles may thus be regulated by differential expression of paraspeckle proteins that have redundant functions in particular cell types in different species. While the domain structures of FET family proteins are conserved between the three proteins, the amino acid compositions of the PrLD are diverse. The N-terminal PrLD is enriched in G, Q, S, Y residues in all FET family proteins, however, a much higher ratio of P and T is observed in EWSR1. The C-terminal PrLDs commonly contain multiple RGG sequences, but an additional septapeptide repeat of YGGDRGG is observed in TAF15. A recent study has shown that FET family proteins undergo a phase transition and form liquid droplets *in vitro* at different protein concentration: FUS formed liquid droplets at the lowest concertation and EWSR1 at the highest concentration among the FET family proteins, and this property was regulated by the amino acid compositions of the N terminal PrLD (Wang et al. 2018). It would be intriguing to study if the dynamics and biophysical properties of paraspeckles are regulated by differential combination of FET family proteins, which may result in cell type-or species-specific biological functions of paraspeckles.

## Materials and Methods

### Animals

All experiments were approved by the safety division of Hokkaido University (#2015-079 and #14-0065) and Kumamoto University (A30-043 and A2020-042). C57BL6/N were used for all experiments for mouse studies. The naked mole-rat colonies were maintained in Hokkaido and Kumamoto University. For anesthetization, medetomidine-midazolam-butorphanol (Kawai et al. 2011) was intraperitoneally injected at a volume of 10 µl/g of body weight.

### Cell culture

NS-Fibroblasts were prepared from adult skin as previously described (Miyawaki et al. 2016). NS-Fibroblasts were cultured in DMEM (high glucose, SIGMA #D5796) at 32°C under humanified atmosphere containing 5% CO_2_ and 5% O_2_. MEF were cultured in DMEM/HamF12 (Wako #044-29765, Japan) at 37°C in 5% CO_2_ and 20% O_2_ conditions.

### Deep sequencing and data analyses

Total RNAs were extracted from NS-Fibroblasts using TRIzol (Thermo Fisher Scientific #15596018). The cellular lysate in TRIzol was incubated at 50°C for 5 minutes to enhance solubilization of semi-extractable lncRNAs as previously described (Chujo et al. 2017). RNA-Seq libraries were made following a standard protocol using Ribo-Zero Gold rRNA Removal Kit (Human/Mouse/Rat) (Illumina) and TruSeq Stranded mRNA Library Prep (Illumina). Deep sequencing was performed using Hiseq4000 in QB3 genomics at UC Berkeley. Sequence reads were mapped to hetGl2 genome using HISAT2 (Kim et al. 2019). To identify n*MALAT1* and n*NEAT1*, the syntenic region of naked mole-rat genome containing *SCYL1* and *FRMD8* was identified by searching the gene names using UCSC genome browser, and mapped reads in the corresponding genomic regions were visualized using IGV genome browser. To estimate the expression of FET family proteins, the number of reads mapped to each gene was counted using featureCounts (Liao et al. 2014) and normalized with the total number of reads and the length of the genes. The sequencing data obtained in this study are available at DRA013388.

### 5’ RACE

5’ RACE was performed using SMARTer RACE 5’/3’ Kit (TAKARA #634858) following the manufacturer’s instructions. cDNAs were synthesized using random primers and used for subsequent adapter ligation and nested PCR to obtain the 5’ end fragments. Obtained PCR products were cloned into a plasmid vector and the sequences were determined following standard procedures. Primers used for the first round of PCR and the second nested PCR are listed in Supplemental table S3.

### Probe preparations

DNA was extracted from the finger of naked mole-rats using a standard protocol. cDNA was synthesized using ReverTra Ace (TOYOBO #TRT-101, Japan) following manufacturer’s instructions. To prepare templates for the probes, PCR was performed using Quick Taq HS DyeMix (TOYOBO #DTM-101, Japan), and genomic DNAs or cDNAs as templates. Primers used to amplify the probe sequences are listed in Supplemental table S3. PCR conditions used was pre-denature at 94°C for 2 minutes, followed by 30 cycles of denature at 94°C for 30 seconds, annealing at 58°C for 30 seconds, and extension at 68°C for 1 minute. Amplified PCR fragments were cloned into pCRII-TOPO using TOPO TA Cloning Kit (Thermo Fisher #452640) following manufacturer’s instructions. To prepare templates for RNA probe synthesis, PCR was performed using the M13 forward and reverse primers and the plasmid DNAs as templates, and the amplified DNA fragments were purified using Wizard SV Gel and PCR Clean-Up System (Promega #A9282). DIG-labelled RNA probes were synthesized using T7 or SP6 RNA polymerase and DIG RNA labeling Mix (Roche #11277073910), and free-nucleotides were removed using CENTRI-SEP Spin Columns (PRINCETON SEPARATIONS #CS-901).

### Northern blot analyses

5 µg of total RNAs were separated on 1% agarose gel containing 2% formaldehyde, and gels were treated with 0.05N NaOH for 20 minutes to enhance transfer of high molecular weight RNAs, which step was essential to detect m*Neat1_2*/n*NEAT1_2*. RNAs were transferred on a positively charged Nylon membrane (Merck #11209299001) using standard protocol, and probe hybridization and detection were performed using DIG Easy hyb (Merck #11603558001), anti-Dig AP (Merck #11093274910), and CSPD-Star (Merck #11685627001) following manufacturer’s instructions. Chemiluminescence signals were detected with ChemiDoc Touch Imaging System (Bio-Rad).

### RT-qPCR analyses

cDNAs were synthesized using ReverTra Ace qPCR RT Master Mix with gDNA Remover (TOYOBO #FSQ-301, Japan) following manufacturer’s instruction. 500 ng of total RNAs were used as templates and 1/50 volumes of synthesized cDNAs were used for subsequent qPCR analyses. qPCR reactions were performed using THUNDERBIRD SYBR qPCR Mix (TOYOBO #QPS-201, Japan) and CFX Connect Real-Time System (Bio-Rad) with the following conditions: 95°C for 1 minutes followed by 40 cycles of 95°C for 15 seconds and 60°C for 60 seconds. The qPCR primers used in this study are listed in Supplemental table S3. Relative expression for each gene was calculated by the ΔCt method using Gapdh as a standard. To obtain relative abundance between FET family mRNAs, the relative expression was further normalized with Ct values obtained by equal amount of PCR fragments (0.1 pmol for each reaction) to normalize the amplification efficiency of primers for each gene.

### *In situ* hybridization

*In situ* hybridization was performed as previously described (Mito et al. 2016). Briefly, cultured cells were fixed in 4% paraformaldehyde in calcium-free Hepes-buffered saline HCMF (10 mM HEPES [pH 7.4], 137 mM NaCl, 5.4 mM KCl, 0.34 mM Na_2_HPO_4_, 5.6 mM glucose) for 10 minutes at room temperature, permeabilized with 0.1% TritonX-100 for 10 minutes at room temperature, and hybridized overnight in 2× SSC containing 50% formamide at 55°C. After washing in 2× SSC containing 50% formamide, probes nonspecifically bound to the cells were removed by RNaseA treatment, and further washed in 2× SSC and 0.2× SSC for 30 minutes at 55°C. Hybridized probes were detected with primary and secondary antibodies listed in Supplemental table S3. To detect mRNAs on tissue sections, tissues were embedded in OCT compounds (Sakura, Japan) and were fresh-frozen in ethanol with dry ice. Cryosections at a thickness of 8 µm were collected on PLL-coated slide glasses, fixed overnight in 4% paraformaldehyde in HCMF. The samples were then treated with 0.2N HCl, 3.3 µg/ml of proteinase K in a 10× TE buffer (100 mM Tris-Cl [pH 8.0] and 10 mM EDTA), and were acetylated with acetic anhydride in triethanolamine buffer before hybridization as previously described (Mito et al. 2016). Probes were detected by chromogenic reaction using alkaline phosphatase-conjugated anti DIG antibody and NBT/BCIP as substrates.

### Production and infection of lentivirus

Lentiviruses were prepared using ViraPower Lentiviral Expression Systems (Thermo Fisher) according to the manufacturer’s instruction. FLAG-tagged full-length cDNA sequences of m*Fus*, m*Ewsr1*, and m*Taf15* were obtained by PCR using primers listed in Supplemental table S3 and cloned into pLenti6/V5-DEST using Gibson Assembly system following the manufacturer’s instruction. For lentiviral infection, m*Fus* KO MEF cells were incubated overnight with culture supernatants obtained from virus-producing 293T cells transfected with packaging vectors, and were further incubated for 48 hours in a fresh medium.

## Acknowledgements

This work was supported by Grants-in-Aid for Scientific Research from JSPS/MEXT granted to S.N. (21H05274, 21K19246, and 17H03604), to K.M. (21H02392 and 21H05143) and to Y.K. (19K06469), and Grant-in-Aid for Early-Career Scientists from JSPS granted to M.K. (21K15012), and JST PRESTO Program (1159399) granted to K.M., and JST FOREST Program (MJFR216C) granted to M.K.

## Figure legends

**Supplemental figure S1.**
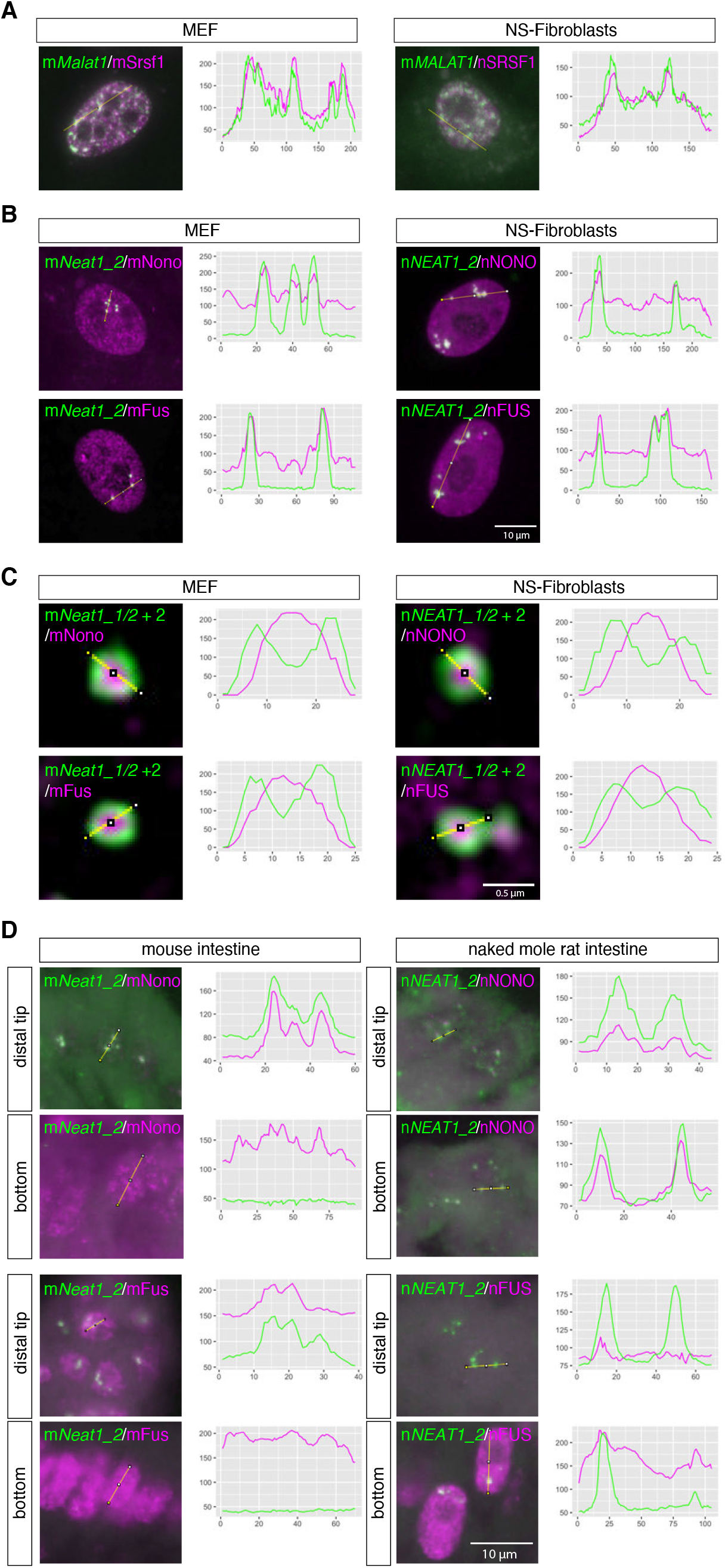
Color profile plots for the images shown in Figure 2A (A), Figure 2B (B), Figure 2C (C), and Figure 4A and 4C (D). Yellow lines indicate the region analyzed in the intensity profile plots. X axis represents the number of pixels and Y axis represents arbitrary unit for the intensity of the signals of each channel. Scale bars, 10 µm in A, B, and D, and 0.5 µm in C.

**Supplemental figure S2.**
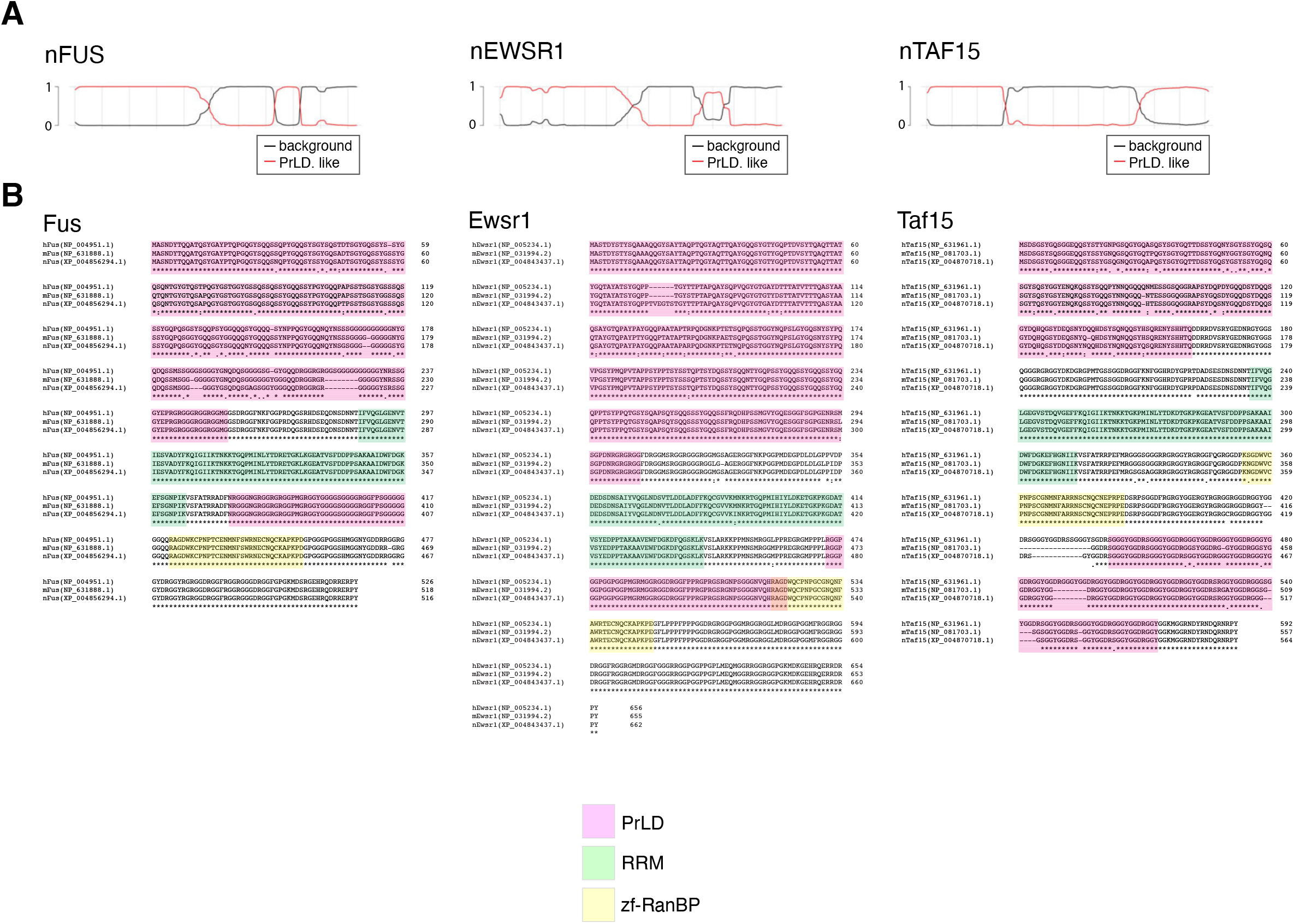
(A) Prion like domains of FET family proteins, predicted by PLAAC (Lancaster et al. 2014). (B) Amino acid sequence alignment of FET family proteins in human (h), mouse (h), and naked mole-rat (n).

**Supplemental Table S3.**
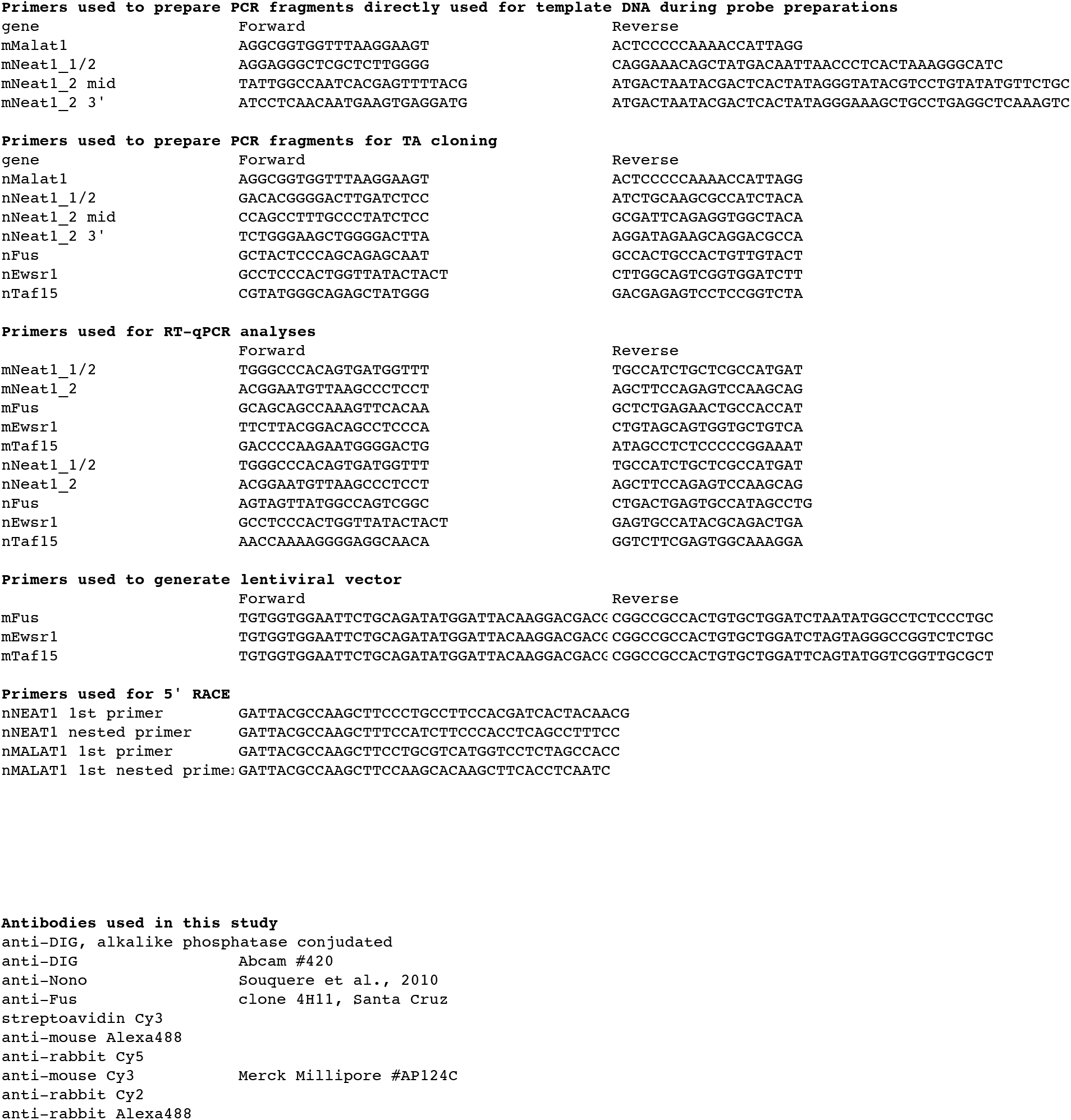
List of antibodies used in this study, and list of primers used to for vector constructions and RT-qPCR analyses.

